# Kit Ligand and Kit receptor tyrosine kinase sustain synaptic inhibition of Purkinje Cells

**DOI:** 10.1101/2023.05.30.542885

**Authors:** Tariq Zaman, Daniel Vogt, Jeremy Prokop, Qusai Abdulkhaliq Alsabia, Gabriel Simms, April Stafford, Bryan W. Luikart, Michael R. Williams

**Affiliations:** Department of Pediatrics & Human Development, College of Human Medicine, Michigan State University; Office of Research, Corewell Health; Department of Molecular and Systems Biology, Geisel School of Medicine at Dartmouth College

**Keywords:** Kit Ligand, Kit Receptor Tyrosine Kinase, Cerebellum, GABAergic Synapse

## Abstract

The cell-type specific expression of ligand/receptor and cell-adhesion molecules is a fundamental mechanism through which neurons regulate connectivity. Here we determine a functional relevance of the long-established mutually exclusive expression of the receptor tyrosine kinase Kit and the trans-membrane protein Kit Ligand by discrete populations of neurons in the mammalian brain. Kit is enriched in molecular layer interneurons (MLIs) of the cerebellar cortex (i.e., stellate and basket cells), while cerebellar Kit Ligand is selectively expressed by a target of their inhibition, Purkinje cells (PCs). By *in vivo* genetic manipulation spanning embryonic development through adulthood, we demonstrate that PC Kit Ligand and MLI Kit are required for, and capable of driving changes in, inhibition of PCs. Collectively, these works in mice demonstrate that the Kit Ligand/Kit receptor dyad sustains mammalian central synapse function and suggest a rationale for the affiliation of Kit mutation with neurodevelopmental disorders.

**Visual Abstract:** 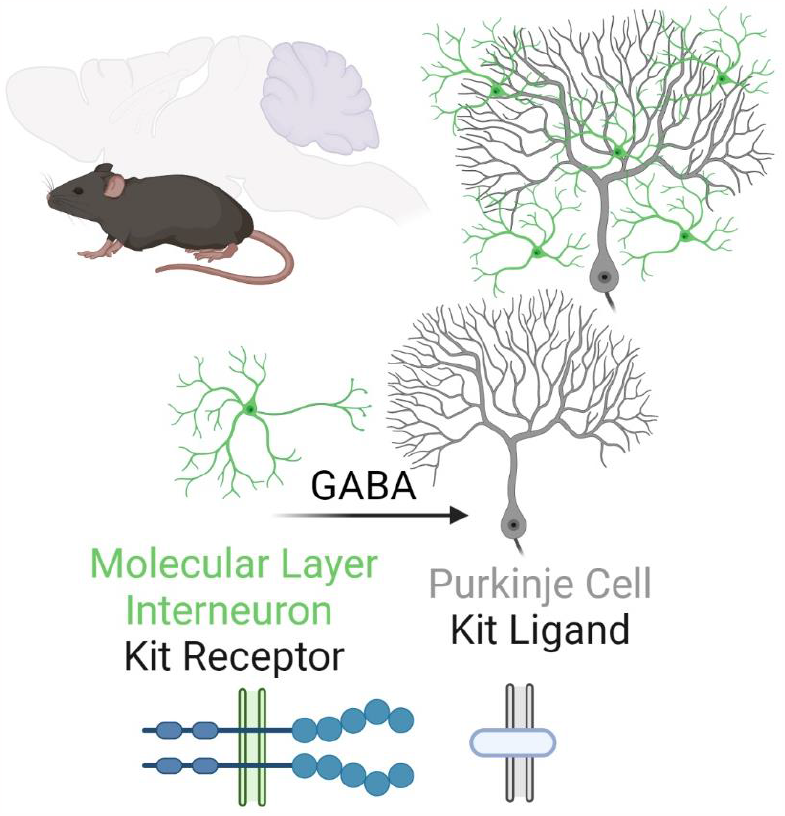

## Introduction

The proto-oncogene receptor tyrosine kinase Kit (c-Kit, CD117) is an evolutionarily conserved, loss-of-function intolerant, gene that is enriched in specific neuron populations and tentatively associated with rare cases of neurological dysfunction (pLI= 0.98 and LOEUF of 0.17) (**Figure S1A,B**) [1, 2]. It is perhaps unsurprising that loss-of-function Kit mutations in humans are rare; Kit is highly pleiotropic, supporting diverse cell populations including melanocytes, hematopoietic stem/progenitor cells, germ cells, masts cell, and interstitial cells of Cajal [3]. Thus, Kit mutations can result in diverse conditions such as hypo- or hyperpigmentation or melanoma, anemia or leukemia, infertility, mastocytosis, and impaired gut motility or gastro-intestinal tumors (Kit biology reviewed [2] [4]).

The activation of Kit is stimulated by Kit Ligand (Gene symbol KITLG, herein KL), a single-pass transmembrane protein having an extracellular active domain that induces Kit dimerization and kinase activity. Across several decades and species, it has been reported that KL and Kit are mutually expressed in discrete neuron populations known to be connected; it has thus been hypothesized that the expression pattern of KL-Kit may reflect a role in connectivity [5-8]. Consistent with such a function, case reports have implicated inactivating Kit mutations with disorders of the central nervous system (References [9-13] and **Figure S1C**). Clinical phenotypes include developmental delay, ataxia, hypotonia, intellectual disability, deafness, and autism spectrum disorder. White matter abnormalities have been noted, and limited evidence suggests KL-Kit may indeed regulate axon outgrowth. KL or Kit mutation or pharmacological manipulation alters outgrowth of spinal commissural axons *in vitro* [14], KL stimulates cortical neurite outgrowth *in vitro* [15], and Kit reduction in cortex of developing rats or mice delays axon extension [16]. While these reports provide anatomical/morphological data suggesting that KL-Kit influences connectivity, these studies did not address consequences to neuronal physiology or synaptic connectivity. Functional studies of KL-Kit in synaptic physiology have likely been hampered by pleiotropy and organismal or cell death. Severe global KL-Kit hypomorphs exhibit fatal anemia, and developmental Kit depletion or inhibition in the cerebrum has resulted in the death of neuronal progenitors or nascent neurons [16-18]. Therefore, whether KL-Kit is necessary for synaptic function has remained largely unstudied. Here, we address this gap.

In mice, rats, and humans, Kit is abundantly expressed by molecular layer interneurons (MLIs) of the cerebellar cortex [6-8, 19], while cerebellar KL is restricted to Purkinje cells (PCs), which MLIs provide GABAergic synaptic inhibition to. We tested the hypothesis that PC KL and MLI Kit were essential to the GABAergic inhibition of PCs. We created a Kit conditional knockout mouse and accomplished embryonic knockout of Kit from MLIs; separately, we knocked out KL from postnatal PCs. By either method, disruption of the KL-Kit pair produced robust and specific impairments to GABAergic inhibition of PCs. Through sparse postnatal viral manipulation of KL in PCs, we provide evidence that KL functions throughout adulthood to regulate synaptic function. These results demonstrate that cell-type specific expression of KL and Kit influences functional connectivity in the mammalian brain, informing the long-observed expression of KL-Kit in the brain and suggesting a rationale for the affiliation of Kit loss with neurological impairment.

## Methods

### Animals

All procedures performed at Michigan State University were approved by the Institutional Animal Care and Use Committees of Michigan State University, which is accredited by the Association for Assessment and Accreditation of Laboratory Animal Care (AAALAC). Mice were on a 12-hour dark/light cycle, with *ad libitum* food and water. Male and female mice were used for all experiments. Mice were genotyped in-house using standard PCR-based methods, or via Transnetyx.

### Creation of Kit conditional knockout mouse

We generated a strain of mice having exon 4 of Kit flanked by LoxP sites (“floxed”). This was accomplished by utilizing the services of UNC Animal Models Core (Dr. Dale Cowley) to generate chimeras from an ES cell line having a Knockout-first Allele (with conditional potential) for Kit (HEPD0509_8). The ES clones were obtained through the EuMMCR (European Mouse Mutant Cell Repository) of the EUCOMM (European Conditional Mouse Mutagenesis Program) [20]. Chimeric mice were screened for germline transmission of the targeted allele and were then crossed to a Flp deleter strain (C57BL/6N-Albino-Rosa26-FlpO) to convert the knockout-first allele to a conditional allele (tm1c) through removal of the FRT-flanked selection cassette. F1 male offspring carrying Kit tm1c and Flp alleles were crossed to C57Bl6/J females; we selected resultant progeny heterozygous for Kit tm1c but negative for Flp and the selection cassette. The Kit tm1c allele was bred to homozygosity while selecting against animals carrying the Tyr c-Brd allele (previously conveyed by C57BL/6N-Albino-Rosa26-FlpO). The resultant strain of mice having the Kit floxed tm1c allele are given the nomenclature C57BL/6N−Kit^tm1c(EUCOMM)Mirow^/J; the Mirow lab code is registered through the National Academies to the author and principal investigator herein, **Mi**chael **Ro**land **W**illiams. Additional strains of mice used were Pax2-Cre transgenic mice (STOCK Tg(Pax2-cre)1Akg/Mmnc, RRID:MMRRC_010569-UNC [21], Kit eGFP (Tg(Kit-EGFP)IF44Gsat/Mmucd; Gensat) [22] backcrossed for greater than 5 generations to C57Bl6/J (Stock No: 000664, The Jackson Laboratory), KitL floxed (Kitltm2.1Sjm/J, Stock #017861) [23], and Pcp2-Cre (Stock # 004146) mice [24], via The Jackson Laboratory.

To produce mixed litters of “Control” and “Kit KO” animals, mice homozygous for the Kit tm1c allele but negative for Pax2-Cre were crossed to mice homozygous for the floxed allele and Pax2-Cre positive. We used Kit KO females crossed to Control males as our breeding scheme. Experimental animals were derived from at least 5 generations of backcrossing Kit KO animals to Kit tm1c homozygous animals derived from a separate (Kit tm1c homozygous x Kit tm1c homozygous) breeding cohort.

### Validation of Kit knockout

We investigated conditional knockout of Kit protein from the cerebellum by Western blot of total protein lysates from acutely harvested cerebellums (∼P52) and by indirect immunofluorescence on free-floating vibratome-generated sections of transcardial-perfused formaldehyde-fixed mouse brains (∼P31). Western blots were probed with Rabbit: α-Kit (1:1000, clone D13A2, Cell Signaling Technologies product 3074), α-GAPDH (1:2000, Clone 14C10, product 2118 Cell Signaling Technology), or α-parvalbumin antibody (1:3000, product PV-27, Swant), each primary was detected via HRP conjugated Goat α-Rabbit secondary antibody (1:4000, Bio-Rad, product 170-6515) after incubation with Clarity ECL substrate (Bio-Rad, 1705061), via a Bio-Rad ChemiDoc MP.

For subsequent imaging and electrophysiology, we focused on lobule IV/V of the cerebellar cortex, as this area was superficially accessible for surgery, and provided a consistent area for focused comparisons between animals that would reduce the influence of spatial variations in KL or Kit expression.

For indirect immunofluorescence, primary antibodies were rat α-Kit (1:500, Rat α-Kit clone ACK4, ThermoFisher, product MA5-17836), mouse α-calbindin (1:2000, α-calbindin-D28K, CB-955, Sigma Aldrich), and rabbit α-parvalbumin (1:500, PV-27, Swant). Primaries were detected by Goat-host highly cross-adsorbed secondary antibodies coupled to Alexa 488, Cy3, or Alexa 647 (1:400, Jackson ImmunoResearch). Nuclei were labeled with DAPI, tissues were mounted in an anti-fade reagent, and fluorescent signals captured laser scanning confocal microscopy. Cell densities for MLIs (parvalbumin positive and/or calbindin negative somas of the molecular layer) were performed on Maximum Z-projections of 2.5 to 2.6 micron thick stacks, using the multi-point feature for counting somas, and the polygon ROI tool for determining the area of the Molecular Layer in which the somas were counted; results from two-technical replicates were averaged for each animal. The polygon tool was also used for determining maximal PSD-95+ pinceau area. To determine the width of the molecular layer, seven linear ROIs per field of view of comparable locations of lobules IV/V were averaged to generate per animal layer thickness measurements. For analysis of synapses, we quantified puncta (of 0.05-5 μm^2^ using the Analyze Particles function) in the molecular layer that were triple positive (each channel auto-threshold) for rabbit α-VGAT(1:500, 131 003), mouse α-Gephyrin (1:5000, 147 011), and Chicken α-Calbindin (1:500, 214 006), all from Synaptic Systems. For analysis of pinceau markers we utilized mouse α-Kv1.1 (1:250, K36/15), Kv1.2 (1:250, K14/16), and PSD-95 (1:200, K28/43), all from NeuroMab/AntibodiesInc.

### Stereotaxic Procedures

Injections of replication-defective viral particles used protocols substantially similar to those previously published [25]. In brief, mice were brought to an anesthetic plane using inhaled isoflurane by a low-flow system (SomnoSuite, Kent); thermal regulation was assisted by heating pad, and pain managed through topical lidocaine and by intraperitoneal ketoprofen. A digital stereotaxic device (Kopf) and reference atlas were used to localize a small hole in the skull; cerebellar coordinates were: neonates; A/P, -2.55; M/L, - 1.1 (reference Lambda), D/V, 1.5 to 0.5 from dura; juvenile; A/P, -6.25; M/L, - 1.35; D/V, 1.5 to 0.75 (reference Bregma); adults; A/P, -6.35; M/L, - 1.8; D/V, 1.5 to 0.75 (reference Bregma). Viral particles were delivered via a ∼30-gauge Hamilton Syringe controlled by digital syringe pump (WPI). The infused virus volume was 1-2 ul and there was a 5-minute period after infusion prior to needle withdrawal. Animals were monitored during and after operations. Ages for injection were, nominally, postnatal day (P) 7, 19, and 56; for P7 and P19, variability of up to two days was utilized to accommodate differences in animal size, and for P56 the eighth week of life was utilized.

### Viruses

Lentiviral particles were derived from our vectors previously described to express transgenes under the hUbiC promoter and pseudotyped with VSV-G [26, 27]. Silent mutations disrupted the internal EcoRI sites of mouse KitL (MC204279, OriGene), the KitL coding sequence was then flanked by EcoRI sites, and the subsequent KitL coding sequence was introduced into a FUC-T2A-(EcoRI) plasmid, to create a FUC-T2A-KL-1 plasmid, wherein C designates mCherry. Subsequent deletion of KitL Exon 6 generated a KL-2 vector [28-31]. Sequences were verified by sequencing. Adeno-Associated Viral (AAV) particles were created by Vigene to drive Cre expression by the Purkinje Cell-Specific L7.6 promoter [32], or to drive mCherry or mCherry-T2A-KL-2 in a Cre-On fashion under the EF1α promoter, with the AAV1 serotype.

### Electrophysiology

For patch clamp electrophysiology of Control vs Kit KO, cells were recorded in slices generated from animals at 36.9 ± 0.6 and 37.4 ± 0.9 days for MLIs and PCs respectively. Analysis of the Pcp2 Cre mediated KL KO tissues was conducted similarly at ∼40-44 days old. For viral PC KL KO animals injected as neonates (P7), juvenile (P19), and adults (P56) were recorded at an average of ∼35, 45, and 83 days old, respectively. Avertin-anesthetized mice were perfused with a carbogen-equilibrated ice-cold slicing solution containing (in mM): 110 C_5_H_14_ClNO, 7 MgCl_2_.6H_2_O, 2.5 KCl, 1.25 NaH_2_PO_4_, 25 NaHCO_3_, 0.5 CaCl_2_-2H_2_O, 10 Glucose, and 1.3 Na-Ascorbate. In the same solution, we generated 250-micron thick parasagittal sections of the cerebellum (Leica VT1200). Slices recovered at 34°C for 30 minutes in a carbogenated ACSF containing (in mM): 125 NaCl, 25 NaHCO_3_, 1.25 NaH_2_PO_4_, 2.5 KCl, 1 MgCl_2_.6H_2_O, 1 CaCl_2_-2H_2_O and 25 Glucose; slices were then held at room temperature for at least 30 minutes before recording. Recordings were performed in carbogenated external recording ACSF (32.7 ± 0.1°C) containing (in mM): 125 NaCl, 25 NaHCO_3_, 1.25 NaH_2_PO_4_, 2.5 KCl, 1 MgCl_2_.6H_2_O, 2 CaCl_2_-2H_2_O and 25 Glucose. MLIs and PCs of folia IV/V were targeted for recording by IR-DIC, or epifluorescence stimulated by a CoolLED pE-4000, detected via a SciCam Pro on a SliceScope Pro 6000-based rig (Scientifica). Recording electrodes were pulled (Narishige, PC-100) from standard-wall borosilicate glass capillary tubing (G150F-4, Warner Instruments) and had 5.1 ± 0.08 and 2.8 ± 0.02 MΩ tip resistance for MLIs and PCs respectively. Spontaneous Inhibitory Postsynaptic Currents (sIPSCs) and Miniature Inhibitory Postsynaptic Currents (mIPSC) were recorded with an intracellular solution containing (in mM):140 CsCl, 4 NaCl, 0.5 CaCl_2_-2H_2_O, 10 HEPES, 5 EGTA, 2 Mg-ATP, and 0.4 Na-GTP, 2 QX-314. For action potential recordings and for Miniature Excitatory Postsynaptic Currents (mEPSCs), the intracellular solution contained (in mM): 140 K-gluconate, 10 KCl, 1 MgCl_2_, 10 HEPES, 0.02 EGTA, 3 Mg-ATP, and 0.5 Na-GTP as described previously [33]. The internal pipette solution pH was adjusted to 7.35 with CsOH for IPSCs and mIPSCs, and with KOH for mEPSCs and action potentials, while for all the osmolarity was adjusted to 300 mOsmolL-1 with sucrose. In whole-cell voltage clamp mode PCs were held at -70 mV; to isolate sIPSCs, CNQX or NBQX (10 μM) and D-AP5 (50 μM) were added to the recording solution and mIPSCs recordings additionally included 1 μM tetrodotoxin (TTX). To record mEPSCs, 1 μM TTX and 50 μM picrotoxin was added to extracellular solution. Action potentials were recorded in MLIs within the middle third of the molecular layer; spontaneous action potentials were recorded in I=0 mode, or action potentials were evoked by depolarizing current injection in 600 ms steps of 10 pA, from 0 to 150 pA, with a 7 second inter-sweep interval. For paired pulse, current ranging from 150 uA to 250 uA was delivered by concentric bipolar stimulating electrode in the molecular layer. Stimulus duration was 50 μS, and ISI was 50 ms. For each cell, 10 sweeps were recorded with 10 S inter-sweep interval. A square-wave voltage stimulation pulse was utilized to determine input resistance and cell capacitance. Purkinje Cells with an access resistance of 10-20 (15.9 ± 0.5), or MLIs with an access resistance <30 (23.6 ± 0.6) MΩ were considered for recording. Recordings with >20% change in series resistance were excluded from analysis. Signals were acquired at 10 KHz with a low-noise data acquisition system (Digidata 1550B) and a Multiclamp700-A amplifier and were analyzed using pClamp11.1 (Molecular Devices) after low pass Bessel (8-pole) filtration (3dB cut off, filter 1 KHz). The minimum amplitude threshold for detecting IPSCs and EPSCs was 15 pA; for mIPSCs and mEPSCs the cutoff was 8 pA. For determining frequency and amplitude, individual events longer than 1 ms were included while overlapping events were manually rejected from analysis. Inhibitory charge is reported as the per-cell sum over the recording epoch of the area under the curve for all pharmacologically resolved inhibitory events. For the analysis of minis, both mIPSCs and mEPSCs with over 8 pA amplitude and longer than 1 ms duration were included while overlapping events were rejected from analysis; also rejected from analysis was any miniature event for which amplitude, rise, and decay metrics were not all available.

### Software

Microscopic image processing was conducted via FIJI/ImageJ, electrophysiological data was analyzed by pClamp11.1, statistical analyses were conducted via GraphPad Prism 9, and Figures were created with BioRender, PowerPoint, and GraphPad Prism 9.

## Results

We generated mice in which Kit exon 4 is flanked by LoxP sites (Kit tm1c); Kit gene modifications are illustrated in **Figure 1C, D**. Kit tm1c homozygous Control animals had normal appearance, sex ratios, and reproductive success. Kit is abundantly expressed in parvalbumin positive GABAergic interneurons in the molecular layer of the cerebellar cortex (MLIs), whereas Kit Ligand (KL) is expressed by Purkinje cells (PCs) (**Figure 1A, B**). To determine if Kit influences synapse function, we conditionally knocked out Kit from MLIs and assessed the MLI:PC synapse. Pax2 is a transcription factor expressed early in the lineage of GABAergic interneurons of the cerebellum [34]; a Pax2-Cre transgene enables embryonic recombination [35, 36]. We generated Kit tm1c homozygous litters of which nominally half were hemizygous for Pax2 Cre (Kit KO) and half were not (Control). We produced Control and Kit KO animals in normal sex and genotype ratios. As expected for the C57Bl6/J background, Control littermates had dark coat and eyes, but littermate Kit KO animals had white whiskers, white fur with variable pigmented patches, and black eyes; example littermates **Figure 1E**. Given Pax2 expression in melanocytes [37] and the role of Kit in melanogenesis (Reviewed in [38, 39]), the pigmentation phenotype provided gross confirmation of conditional Kit KO. We thus confirmed that cerebella of Kit KO animals were Kit depleted. By immunohistochemistry in Control animals, we recapitulated the known pattern of Kit immunoreactivity in the parvalbumin positive MLIs of the cerebellar cortex (Control, **1F and Inset**). In contrast, a sex-matched Kit KO littermate demonstrated loss of cerebellar Kit immunoreactivity (Kit KO, **1F and Inset**). In data not shown, we confirmed that both mGlur1/2 and neurogranin positive cerebellar Golgi cells (which share the MLI Pax2+ lineage, and of which some are transiently Kit positive) were still present in the Kit KO condition [34, 40-42]. By Western blot of cerebellar lysates from Control animals, Kit resolved as a major band at ∼120 kD, but in Kit KO littermates, Kit immunoreactivity was lost; GAPDH as loading control was equivalent, as was parvalbumin (which marks mature MLIs and PCs) **Figure 1G**.

**Figure 1:**
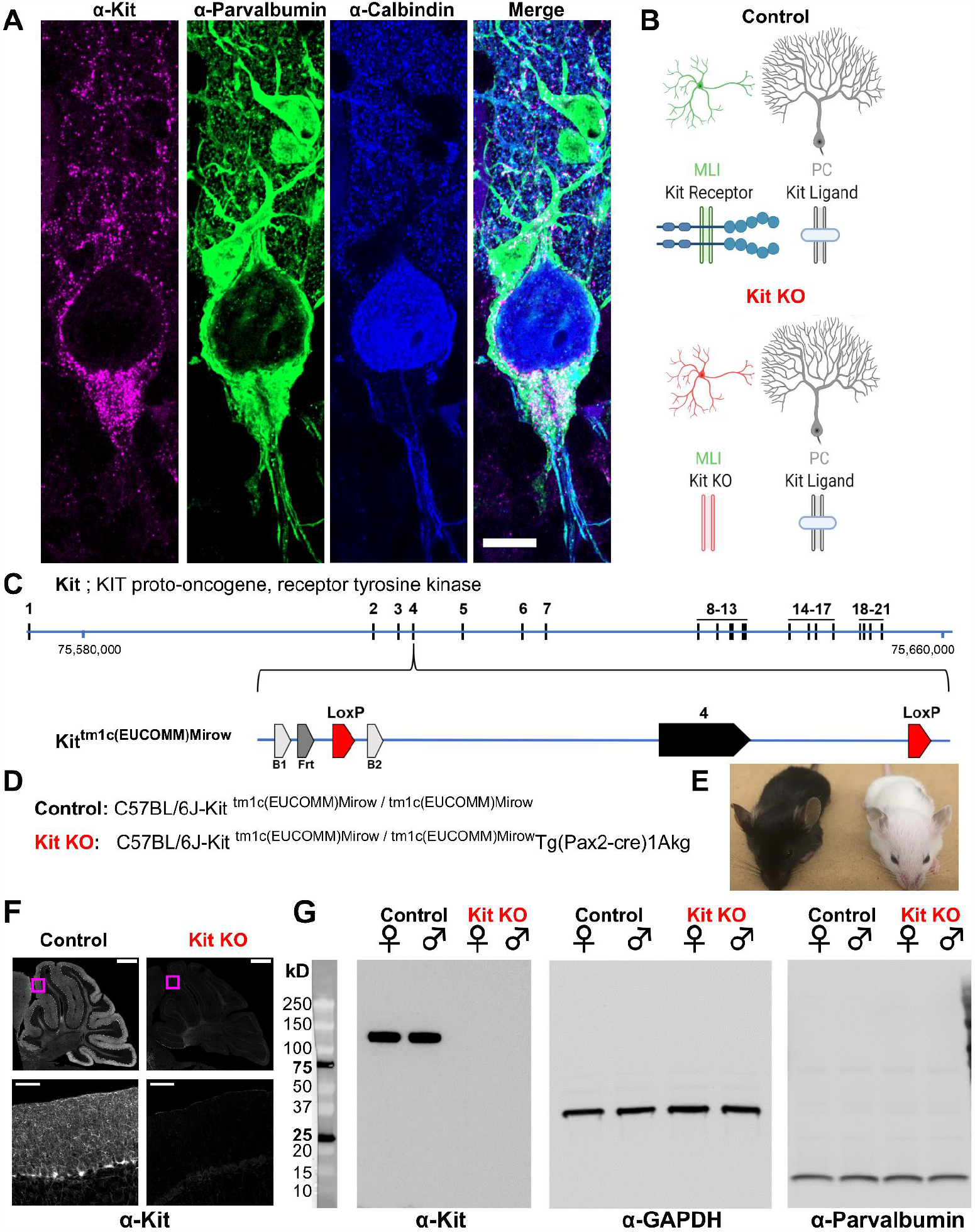
Design and validation of a Kit conditional knockout mouse. (**A, B**) Kit receptor tyrosine kinase (Kit) is enriched in parvalbumin positive GABAergic interneurons of the molecular layer (i.e. basket and stellate cells, MLIs) of the cerebellar cortex, where they synapse onto each other and onto Purkinje cells (PCs, Calbindin+), which express Kit Ligand (KL). Scale bar 10 microns. Expression pattern schematized in B, Control. **(C)**In humans and mice, Kit is encoded by up to 21 exons, which in mouse is encoded on the plus strand of chromosome 5 at 75,735,647-75,817,382 bp. We generated a Kit conditional knockout mouse in which Kit Exon 4 is floxed, flanked by LoxP sites. **(D)**We generated Control mice homozygous for the Kit floxed allele Kit^tm1c(EUCOMM)Mirow^, which varied in Pax2-Cre)1Akg transgene status, with the goal of depleting Kit from MLIs in embryonic development. **(E)**Pax2 Cre mediated Kit KO mice were notably hypopigmented in hair and whiskers, though not eyes. **(F)**Confocal microscopy of Kit immunoreactivity in cerebella from age and sex matched Control and Kit KO littermates demonstrates the established enrichment of Kit in the molecular layer of the Control cerebellar cortex, and its loss in Kit KO. Scale Bar 500 microns top row, 50 microns inset. **(G)**Utilizing a distinct assay and different primary antibody, we confirm the detection of Kit immunoreactivity in Controls, and its loss in Kit KO litter mates of either sex by Western Blot of total protein lysates of cerebella. We affirmed equivalent protein loading by GAPDH and parvalbumin.

Having validated that Kit KO animals indeed lacked cerebellar Kit, we then assessed the functional impact. We compared spontaneous GABAergic inhibitory post synaptic currents (sIPSC) in PCs of acute slices generated from ∼P34 Control and Kit KO animals, schema **Figure 2A**, example traces **Figure 2B**. Kit KO was associated with a significant change in the distribution of sIPSC event amplitudes (**Figure 2C**). For each PC of Control or Kit KO conditions, we calculated the average sIPSC event frequency and amplitude, as well as total inhibitory charge transfer over the recording epoch. We determined there was a significant decrease in sIPSC frequency (**Figure 2D**) and a less severe decrease of sIPSC amplitude and inhibitory charge. This reduced inhibition did not produce obvious impacts to spontaneous PC firing dynamics: we determined (in 29 Control vs 22 Kit KO cells) that PC membrane potential and membrane resistivity were not significantly different, and that per cell average: spontaneous action potential frequency, inter-spike interval (ISI), and ISI CV2 were not significantly different, despite some difference in the distribution of individual ISI across genotypes.(**Figure S6**). Though circuit consequences of the Kit KO mediated PC disinhibition were thus not immediately apparent, the reduced sIPSC frequency in Kit KO was not associated with changes in the density or distribution of MLIs or in the thickness of the molecular layer (**Figure S2)**.

**Figure 2:**
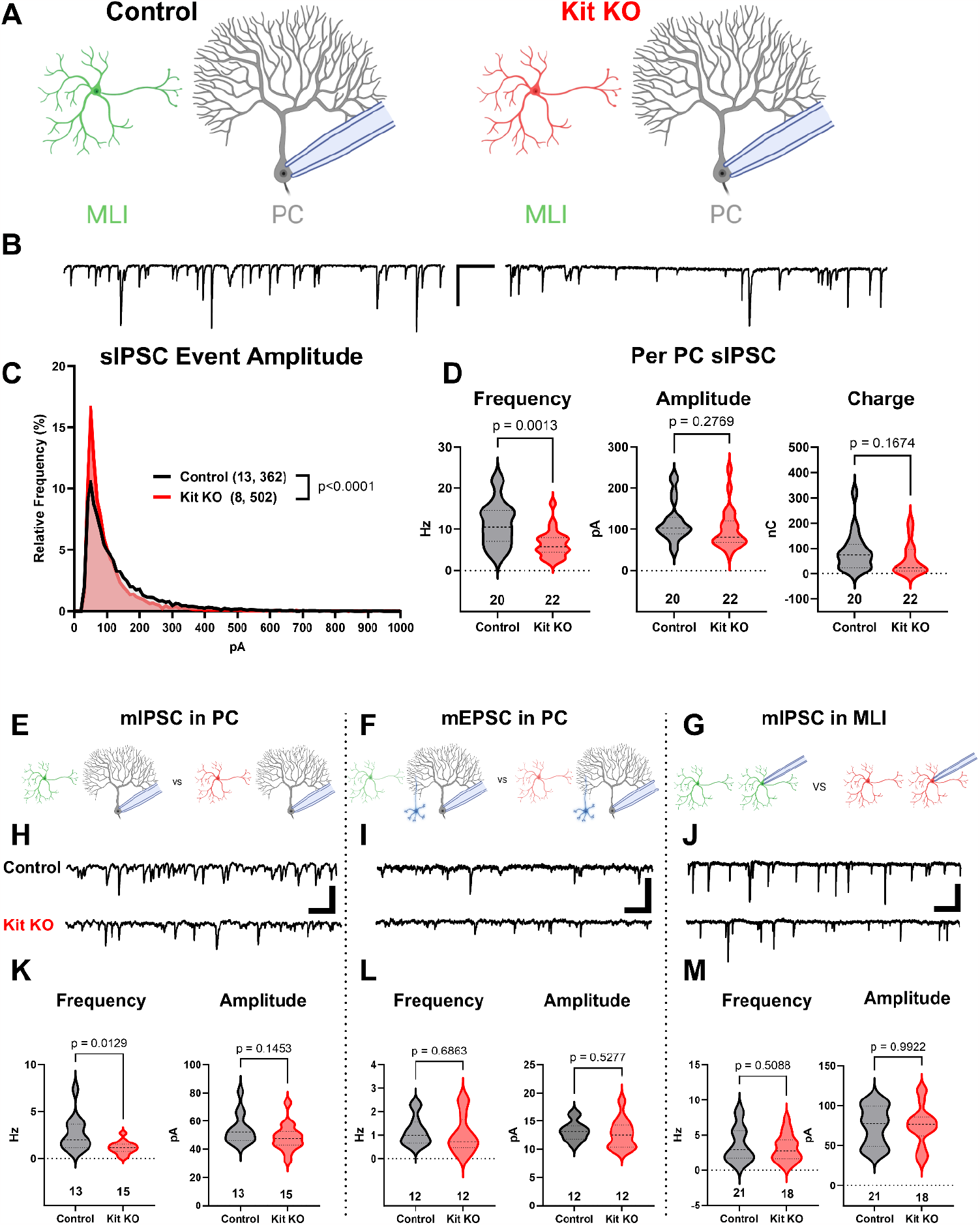
The knockout of Kit from Cerebellar Cortex Interneurons Impairs Inhibition of Purkinje Cells. 419. (**A, B**) Experimental schema and example traces of spontaneous inhibitory postsynaptic currents in Purkinje Cells (PCs) from Control animals or from those with Kit KO from cerebellar cortex molecular layer interneurons (MLIs). Scale bar is 500 ms x 100 pA. **(C)**A frequency distribution plot for individual sIPSC event amplitudes recorded in PCs as in A,B. A KS test reveals a significant difference in the distribution of these event amplitudes; p<0.0001, n in chart. **(D)**For each PC, the average sIPSC Frequency and Amplitude, and the total Inhibitory Charge transfer, was determined. There was a significant ∼50% decrease in sIPSC frequency, but the decrease in Amplitude or Charge transfer was not significant. (**E-G**) Experimental schema: In separate experiments, we recorded miniature postsynaptic currents from PCs or from MLIs in Control and in Kit KO animals. (**H**) Example traces of mIPSCs or (I) mEPSCs recorded in PCs, and example traces of mIPSC in MLIs, all from Control or from Kit KO animals. Scale for H and J is 50 pA, 25 pA for I, all 500 ms. (**K-M**) Analysis of per cell average miniature event Frequency and Amplitude revealed that Kit KO significantly reduced mIPSC frequency, but not amplitude, in PCs by >50%, p=0.013. mEPSCs in PCs and mIPSC in MLIs were not significantly different between Control and Kit KO. n in charts, refers to the number of cells. Error bars are SEM. p-values were calculated by a two-tailed t-test, with Welch’s correction as needed.

Since the number and distribution of MLIs were normal in Kit KO, we sought to determine if the reduced PC inhibition was related to altered MLI physiology (Schema, **Figure S3A**). However, capacitance, input resistance, and spontaneous or evoked MLI action potential frequency was not different between Control and Kit KO (**Figure S3B-E)**. Since MLIs numbers, distribution, and firing were apparently normal in Kit KO, we sought to determine if there were defects in evoked neurotransmitter release. Direct electrode stimulation of the outer or inner molecular layer in a paired-pulse paradigm (Schema, **Figure S3F,J**; example traces **Figure G,K**) evoked IPSCs of equivalent mean amplitude in PCs from Control and Kit KO (**Figure S3H,L**), and we detected no difference in the paired pulse ratio between conditions (**Figure S3I,M**). These data suggest that direct stimulation can evoke a similar maximum of release probability between Control and Kit KO conditions. We therefore investigated markers of the MLI:PC synapses to determine if they were altered.

The density and average size of GABAergic synapses onto PCs (triple positive VGAT, Gephyrin, and Calbindin puncta in the molecular layer) was not reduced by Kit KO (example immunofluorescence, **Figure S4A**, Quantification **S4B,C**). MLI axon terminals contribute not just to individual axo-dendritic synaptic puncta, but also to interdigitated MLI axon collaterals around the soma and initial axon segment of PCs, forming “pinceaux” structures. By parvalbumin immunoreactivity, these structures were preserved but smaller in Kit KO (arrowheads **Figure S4C)**, and this was confirmed by decreased Kv1.2 (**Figure S4D**) and PSD-95 (**Figure S4E**). As PSD-95 immunoreactivity faithfully follows multiple markers of pinceaux size [43], we quantified PSD-95 immunoreactive pinceau area and we determined that pinceaux size was decreased by ∼50% in Kit KO (26 Control vs 43 Kit KO pinceau from n of 5 Control vs 8 Kit KO animals, **Figure S4F**). These results suggested that Kit KO MLIs may have defects in physical and/or functional maturation of synapses onto PCs.

To evaluate the number of functional GABAergic synapses, we analyzed miniature synaptic currents in acute cerebellar slices, (schema **Figure 2 E, F, G**; example traces **Figure 2 H, I, J)**. In PCs of Kit KO, there was a significant ∼50% reduction in the average frequency, but not amplitude, of mIPSCs (**Figure 2K**). In contrast, the average frequency and amplitude of mEPSCs in PCs from Control and Kit KO did not differ (**Figure 2L**). These data suggested that Kit KO impaired the MLI:PC synapse but did not rule out a general defect in MLI synapse function. As MLIs synapse not just onto PCs, but also onto other MLIs, we evaluated mIPSCs in MLIs. Mean mIPSC frequency and amplitude in MLIs was unchanged between Control and Kit KO (**Figure 2M**). Therefore, embryonic MLI Kit KO was associated with a specific defect in the number or proportion of functional GABAergic synapses onto PCs. As MLI Kit and PC KL expression is maintained postnatally, we hypothesized that postnatal postsynaptic PC KL KO would phenocopy embryonic presynaptic MLI Kit KO.

We utilized previously described KL floxed [23] and Pcp2-Cre [24] strains to yield mice with Control or KL KO PCs (Schema **Figure 3A**). We confirmed depletion of cerebellar KL transcripts by rtPCR in data not shown[23], and we confirmed that PC KL KO did not grossly disrupt the pattern of Kit immunoreactivity. Patch clamp of PCs in acute slices revealed that PC KL KO produced significant differences in the distribution of sIPSC event amplitudes (Example traces **Figure 3B, Figure 3C**). PC KL KO reduced the average sIPSC frequency and amplitude, as well as the total inhibitory charge transfer (“Charge”) recorded in PCs (**Figure 3D**), without changes to PC capacitance or membrane resistance. The phenotype was specific to GABAergic synapses: there was a substantial reduction in the frequency but not the amplitude of mIPSC events in PCs (**Figure 3E**). As with Kit KO, we detected no decrease in the per-animal average size or density of triple positive VGAT/Gephyrin/Calbindin synaptic puncta in the molecular layer (n=4 Control vs 4 KL KO animals, mean area 0.249 vs 0.286 μm^2^, p=.1962; mean density 0.045 vs 0.046 per μm^2^, p=.9658), Not Illustrated). We further detected no difference either in PDS-95+ pinceau area (49 Control vs 48 KL KO pinceaux from n=4 Control vs 4 KL KO animals, mean area 57.93 vs 53.37 μm^2^ p=.6036, Not Illustrated). Thus, as with Kit embryonic Kit KO, postnatal KL KO apparently reduced the proportion of functional, if not the absolute structural total, of GABAergic synapse sites upon PCs. In contrast to this impact on GABAergic synapses, the frequency and amplitude of mEPSCs in KL KO PCs was not different vs Controls (**Figure 3F**). In data not shown, a KS test revealed a significant difference in the individual mEPSC event amplitude distribution (p=0.0002, n=1196 Control vs 1244 KL KO), however the mean mEPSC amplitude was within 1% between conditions. Similarly, a KS test of mIPSC event amplitudes revealed that the mean mIPSC amplitude was ∼10% greater in PC KL KO than in Control (p<0.0001, n=5996 Control vs 3958 KL KO). It is therefore unlikely that decreased event amplitude contributed to the specific and robust >50% decrease in mIPSC frequency detected in PC KL KO. Thus, as with MLI Kit KO, PC KL KO specifically reduced synaptic inhibition of PCs.

**Figure 3:**
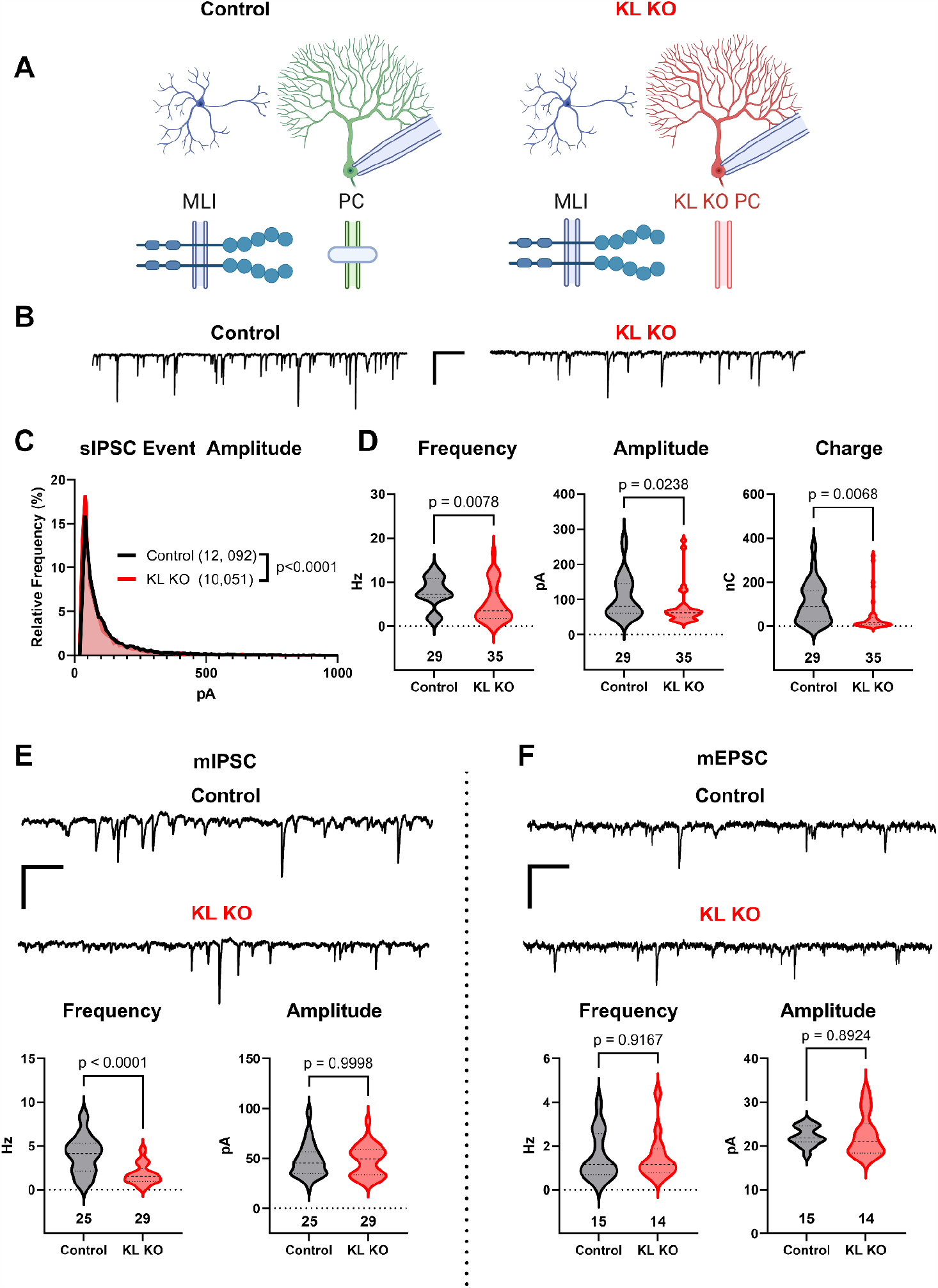
Knockout of Kit Ligand from Purkinje Cells decreases the inhibitory input they receive. (**A, B**) Experimental schema and example traces of inhibitory postsynaptic currents detected in Purkinje Cells (PCs) from Control animals or from those with Kit Ligand knockout (KL KO) accomplished by a Pcp2-Cre x KL floxed strategy. Scale bar is 500 ms x 100 pA. **C**) A frequency distribution plot for individual sIPSC event amplitudes recorded in PCs as in **A, B**. A KS test reveals a significant difference in the distribution of these event amplitudes; p<0.0001, n in chart. (**D**) For each PC, the average sIPSC Frequency and Amplitude, and the total inhibitory Charge transfer, was determined. There was a significant ∼40% decrease in sIPSC Frequency, Amplitude, and Charge transferred to KL KO PCs. (**E, F**) The average mIPSC recorded in PC KL KO vs Control PCs had a >50% decreased Frequency p<0.0001 without a corresponding decrease in Amplitude. The average mEPSC Frequency and Amplitude recorded in separate PCs did not differ between Control and KL KO. Scale bar is 500 ms x 100 pA for mIPSC, 500 ms x 50 pA for mEPSC. n in charts refers to number of cells. Error bars are SEM. p-values were calculated by a two-tailed t-test, with Welch’s correction as needed.

Recombination by Pcp2 Cre begins at ∼P7 [24], whereas Pax2 Cre recombination occurs at ∼E15 [21, 34]. Our results thus suggest that continued expression of the KL-Kit dyad sustains inhibitory drive to PCs. To test this hypothesis (and to avoid transformative effects of Kit overexpression), we tuned KL postnatal expression *in vivo*.

To determine if acute focal manipulations in KL modulate PC inhibition, we injected replication-defective viruses leveraging the PC-specific L7.6 promoter [32] to express Cre in KL floxed animals, and we identified Cre-positive KL KO PCs by mCherry expression (Schema **Figure 4A**, Example transduction, **Figure S5A-C,E-G**). Injections were at P7, P18, or P56; acute slices were generated 2-3 weeks after injections. In the P7 neonatal animals we utilized lentivirus owing to its reduced spread vs AAV particles, which were used for P18 and P56. Patch-clamp recordings in areas of sparse infection revealed that regardless of when KL was depleted, PC KL KO reduced PC sIPSC frequency and inhibitory charge transferred (P7, P18, and P56 in **Figure S5 D, H, I;** P18 injections **Figure 4 B, C, D)**. We next examined the effects of PC KL over-expression using a complementary strategy. We co-injected L7.6 Cre AAV with AAV encoding EF1α driven Cre-on mCherry and the membrane bound isoform of mouse KL (mKL2) via a T2A element to produce PC KL overexpression (OX) (Schema **Figure 4E**). Whether at P18 (shown) or P56 (not shown), KL OX PCs had increased sIPSC frequency, amplitude, and inhibitory charge transferred, compared to nearby Control PCs (**Figure 4F, G, H**). We noted that Control PCs from the PC KL KO paradigm received a similar degree of inhibition (frequency, amplitude, charge) as did PCs from Sham injections. In contrast, while the KL OX PCs received elevated GABAergic input compared to their adjacent Control PCs, the (KL OX) relative local increase in inhibitory drive was not beyond the level of inhibition recorded in PCs from separate Sham animals (**Figure 4F, G, H**). That is, the local gain in synaptic input to KL OX PCs appears to come at the expense of decreasing inhibition to nearby Control PCs.

**Figure 4.**
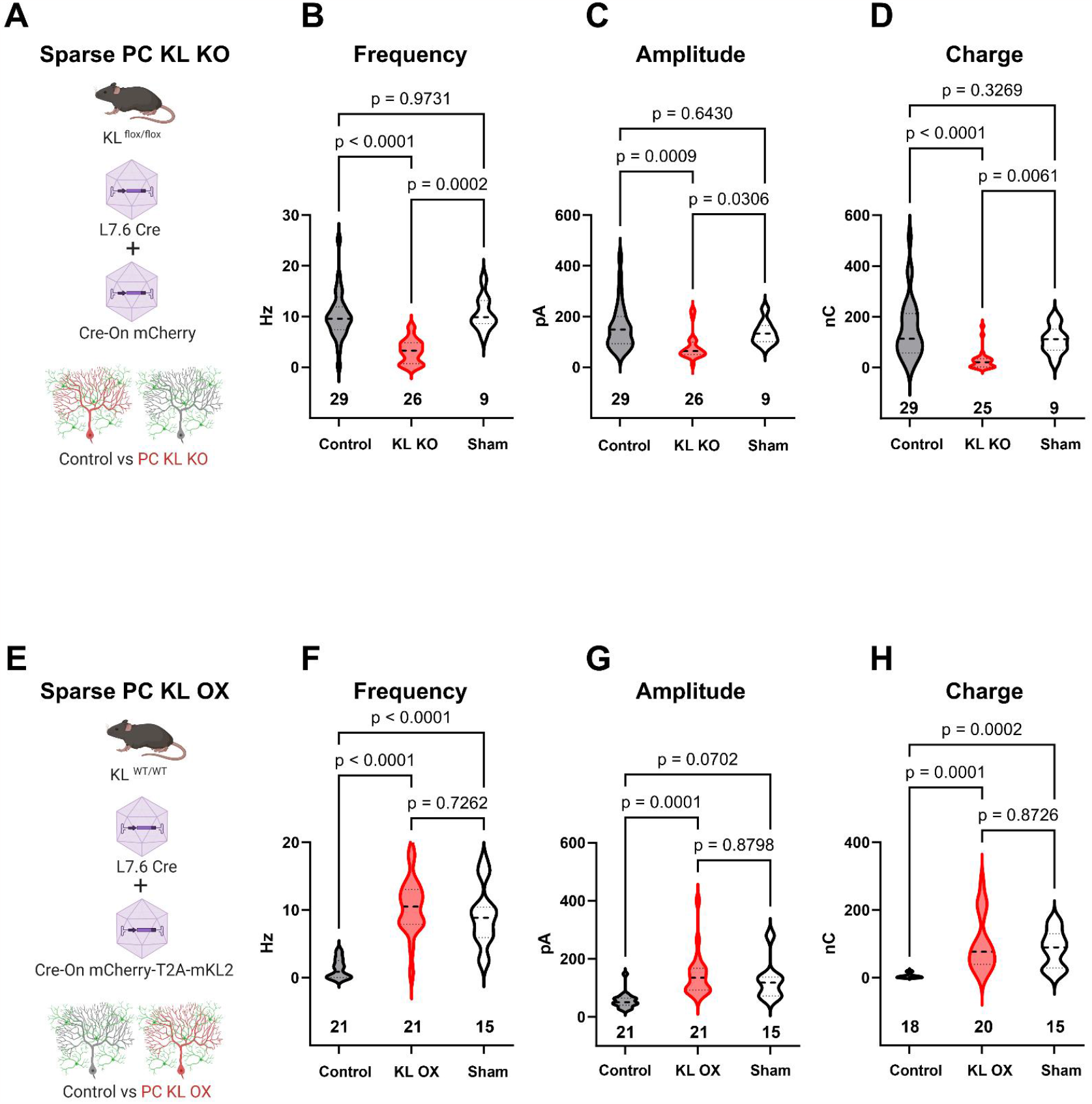
Local levels of Kit Ligand influence the inhibition Purkinje cells receive. (**A**) To accomplish *in vivo* Control and sparse Kit Ligand Knockout (KL KO) Purkinje Cells (PCs), an AAV encoding Cre under the PC specific L7.6 promoter was co-injected with an AAV encoding a Cre-On mCherry cassette under the Ef1α promoter. (**B, C, D**) In animals injected at P18, PC KL KO neurons demonstrated a ∼70% decrease in sIPSC event frequency compared to adjacent Control PCs (p<0.0001) or PCs recorded from Sham control animals (p=0.0002). Control vs Sham sIPSC frequency was not significantly different. The same pattern was found for both mean sIPSC Amplitude (C) and for total inhibitory Charge transferred (D). **(E)**To accomplish *in vivo* Control and sparse Kit Ligand overexpressing (KL OX) PCs, the L7.6 Cre AAV was co-injected with AAV expressing mCherry and (T2A) murine Kit Ligand isoform 2 under the Ef1α promoter. (**F, G, H**) In animals injected at P18, PC KL OX neurons demonstrated an ∼8-fold increase in sIPSC Frequency vs neighboring Control PCs (p<0.0001). The frequency of sIPSC events in KL OX PCs was not significantly different from PCs recorded in different Sham animals; this Sham PC sIPSC frequency (while comparable across studies) was 7-fold higher than Control PCs within the KL OX experimental animals (p<0.0001). A similar pattern was found for sIPSC amplitude (G) and for total inhibitory Charge transferred (H). n in charts, refers to number of cells. Error bars are SEM. p-values were calculated by Brown-Forsythe ANOVA test with Dunnett’s T3 multiple comparisons test.

## Discussion

For genes essential to survival or reproduction, germline knockout strategies are not feasible for evaluating postnatal physiology. Among such genes, Kit is an example of a highly pleiotropic gene, and so the interpretation of global hypomorphs is challenging and the recombination of conditional alleles outside of desired tissues can lead to lethal or sterile phenotypes. By generating a conditional knockout mouse, we were able to produce viable animals with Kit depleted from the cerebellum to reveal a role in synapse function.

MLI number, distribution, action potential firing, and the density of GABAergic synaptic puncta onto PCs, are all apparently normal in MLI Kit KO. These data suggest that while MLI axon terminals are present in Kit KO, they may be less functional. An increased failure rate between action potential generation and neurotransmitter release would be consistent with the normal firing frequency and synapse puncta density but decreased sIPSC and mIPSC frequency observed. Under the PPR paradigm, the apparently equivalent evoked responses may be due to direct stimulation of axon terminals and/or simultaneous recruitment of multiple MLIs.

Kit KO most directly impacts the MLI:PC synapse, based on seemingly normal miniature events for parallel fiber: PC synapses (mEPSC in PC) and MLI:MLI (mIPSC in MLI) synapses. This is perhaps expected given the cell type specific expression of KL by PCs and Kit by MLIs, however, it is a critical distinction as Cerebellins [44] and Calsystenin [45] impact multiple forms of synaptic input to PCs. It will be informative to determine if the residual inhibitory drive to PCs observed in MLI Kit KO or PC KL KO comes from PC axon collaterals [46-49] whose input we might expect to be preserved if KL-Kit functions primarily to mediate connectivity between different cell types, or perhaps from a subtype of interneuron not dependent upon Kit signaling. This distinction is further relevant since our histological analysis of synaptic puncta does not resolve between MLI:PC and PC:PC synapses, the relative balance of which may be altered by either KL or Kit manipulations. Indeed, future studies of the role of KL-Kit on PC-PC connectivity may inform how PC KL OX can apparently weaken inhibition to adjacent PCs: this may occur through the reduction of inhibitory drive from KL OX PCs to adjacent control PCs. An alternative, and not mutually exclusive possibility is that PC KL OX excessively potentiates MLI Kit to enhance MLI:MLI inhibition, reducing the inhibitory output surrounding MLIs can provide to control PCs.

Though such important nuances of the circuit mechanisms remain to be elucidated, our results demonstrate that the KL-Kit axis, from embryonic development through young adulthood, can alter GABAergic drive to PCs. These results suggest that KL-Kit may be necessary to maintain, and perhaps capable of modulating, MLI:PC GABAergic inhibition in the adult cerebellum. As each MLI (e.g. basket cell) has axon collaterals that can target multiple PCs, and as each PC can receive input from multiple MLIs, it is interesting to speculate that dynamics (e.g. in the expression or shedding) of KL may act through presynaptic Kit to tune the degree of convergent MLI:PC inhibitory drive. This would complement existing forms of modulation at the MLI:PC synapse, such as those dependent on other PC derived signaling molecules, like endocannabinoids [50-52] or peptides like secretin [53, 54] and CR [55]. The physiology of PCs, and the structure of MLI:PC inhibition, also varies over zones such as those identified by Zebrin-II and PLC β4 [56-60]; whether there is functional variation in KL or Kit expression, signaling, or dependency, remains to be determined. Though we have demonstrated a role in synaptic GABAergic inhibition, it is currently unknown whether Kit signaling also plays a role in other forms of inhibition in the cerebellar cortex, such as MLI:MLI gap-junctions, or MLI:PC ephaptic coupling. The latter may be suggested by the Kit KO mediated reduction in Kv1.1/Kv.12 and PSD-95 immunoreactivity.

While KL-Kit clearly influences synaptic function, we do not yet know the molecular mechanisms involved. Kit is a receptor tyrosine kinase that can activate multiple downstream effectors including Src family kinases, PLC, PI3K/mTOR, and the MAPK pathway, the latter two notable here for their shared affiliation with synapse phenotypes and autism spectrum disorder. Investigation of the interactome of Kit under basal and KL manipulated conditions may inform the receptor kinase cascades through which Kit signaling sustains synaptic function. It also plausible that trans-cellular interactions between KL and Kit may function in a kinase-independent, cell adhesion molecule modality [61, 62]. There is a substantial literature illustrating the importance of adhesion molecules in the organization and maintenance of MLI:PC inhibition, such as neurexins/neuroligins, dystrophin/dystroglycan, semaphorins, and neurofascins [44, 45, 63-67]. Much of what is understood about these molecules in the maintenance of PC inhibition is revealed by post-synaptic phenotypes; that is, cis-and trans interacting proteins that, via gephyrin, maintain the organization of PC GABA_A_ receptors. It is notable therefore that MLI Kit or PC KL KO has a more predominant effect on mIPSC frequency than on amplitude, suggesting a presynaptic phenotype.

Despite the reduced PC inhibition resulting from MLI Kit KO or from PC KL KO, we have not observed overt cerebellar signs (i.e., ataxia). We suspect that this is due to compensation. Even the developmental ablation of functional GABA_A_ channels in PCs does not produce overt motor signs, while adulthood ablation can [68]. Thus, we might unmask behavioral phenotypes if KL or Kit KO is delayed until adulthood. Consistent with the interpretation that developmental compensation may be involved, the impact of postnatal PC KL KO on inhibitory drive to PCs is more pronounced than embryonic MLI Kit KO. Beyond direct inhibition of PCs, MLI:PC GABAergic inhibition constrains granule cell mediated PC dendritic excitation and thus gates the conversion of PC LTD to LTP at the PF-PC synapse in motor learning [69, 70]. It is therefore possible that KL or Kit KO may only reveal phenotypes in motor learning (or other cerebellar-plasticity dependent) behaviors; speaking to the feasibility of this possibility, global hypomorphs for KL or Kit have altered hippocampal neurophysiology and learning performance on a Water Maze task [71-73]. Continuous expression of KL and Kit in specific neurons may thus reflect not only a role in the development (or maintenance) of synaptic function as suggested here, but also in the plastic modulation of synaptic strength.

Despite the importance of future works to determine molecular mechanisms and organismal consequences, our study is a discrete advance because it establishes that KL-Kit regulates mammalian central synapse function. Here, we demonstrate impacts at the mouse GABAergic MLI:PC synapse; however, Kit and KL exist in diverse organisms and neuronal populations including glutamatergic neurons of the hippocampus, and neurons in olfactory bulb, basal forebrain, and brainstem. It is therefore feasible that the synaptic phenotypes reported here reflect that KL-Kit is broadly capable of influencing connectivity.

## Supplementary Information

### Six Figures and Associated Legends

**Figure S1:**
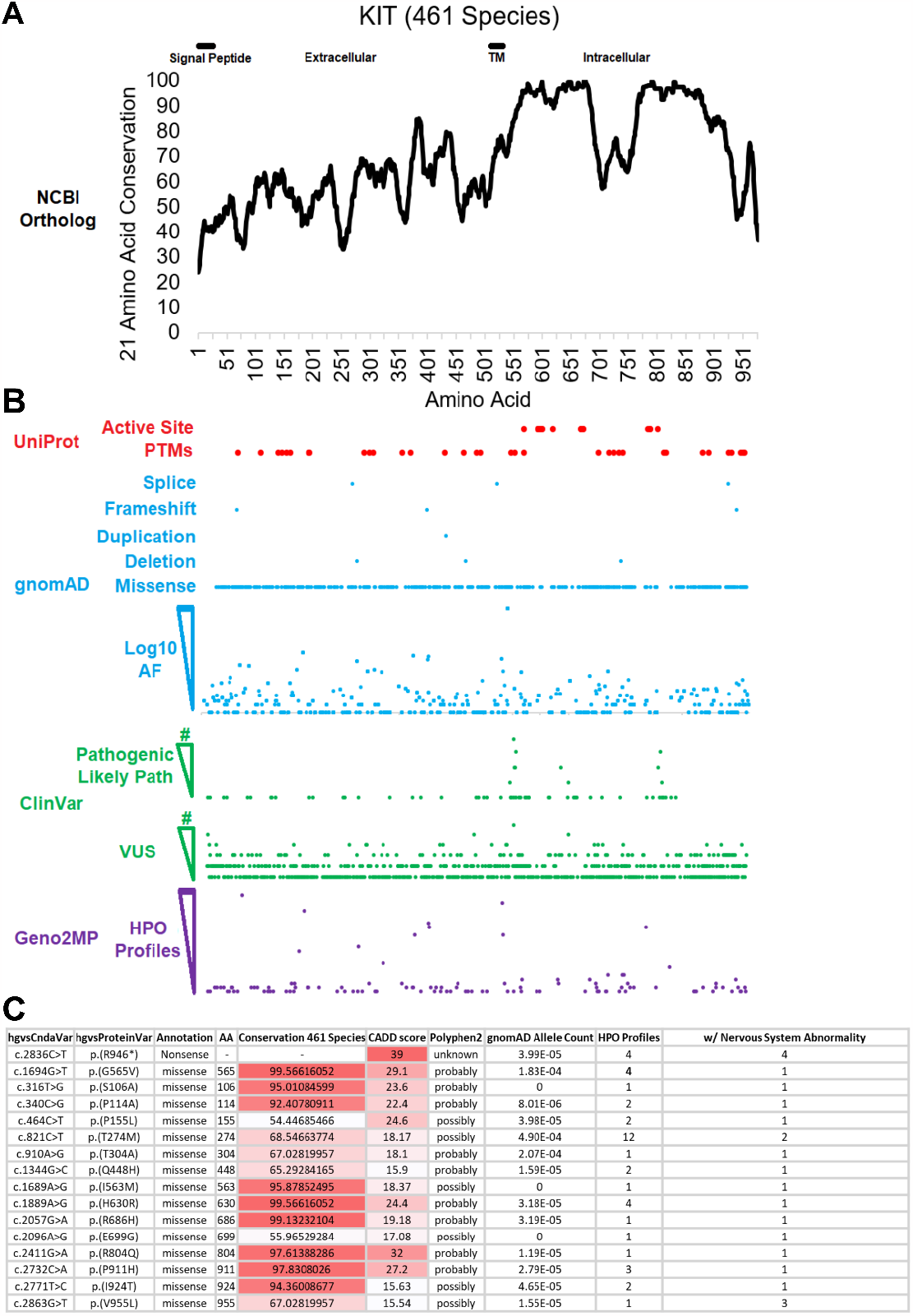
Kit receptor tyrosine kinase is a highly conserved gene affiliated with neurological impairment. **(A)**KIT protein sequences were extracted from NCBI ortholog on March 2023 and aligned with MUSCLE. Conservation was placed on a 21 amino acid sliding window to calculate linear motifs. The intracellular domain possesses a split cytoplasmic kinase motif demonstrating high conservation across 461 species. **(B)**Alignment of Active Site and Post-Translational Motif (PTMs) to Kit amino acid conservation and gnomAD, ClinVar, and Geno2MP human variants. **(C)**Human Genome Variation Society annotations for human DNA and corresponding encoded amino acid changes affiliated with their mutational class (Annotation), Site (AA), and Conservation across 461 species. Combined Annotation Dependent Depletion (CADD) score for deleteriousness of SNV or indel variants. Polymorphism Phenotyping v2 (PolyPhen2) annotations predict impact of amino acid substitution on Kit structure function. Allele counts in Gnomad and the number of profiles in the Human Phenotype Ontology associated with Nervous System Abnormality.

**Figure S2:**
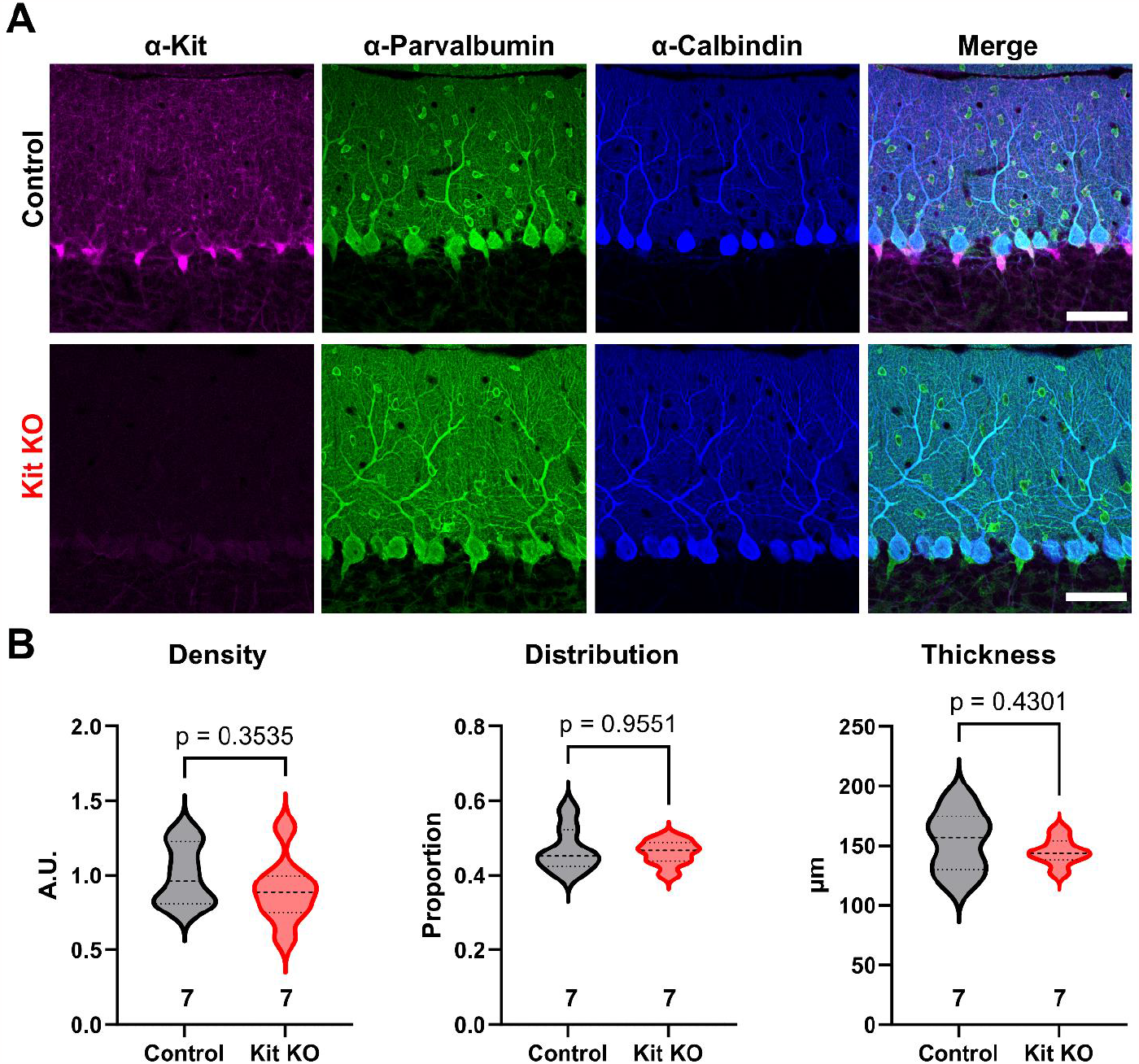
Kit knockout does not alter molecular layer interneuron number or distribution. **(A)**Immunohistochemistry and confocal microscopy of Control and Kit KO mouse cerebellar cortex validates the normal enrichment and targeted depletion of Kit in Control and Kit KO animals respectively. Parvalbumin immunoreactivity is present in both Molecular Layer Interneurons and Purkinje Cells, while the latter is selectively Calbindin immunoreactive. Scale bar 50 microns. **(B)**Quantification of the density of Parvalbumin positive and or Calbindin negative somas within the molecular layer reveals that the Density of MLIs is not reduced by Kit KO. Quantification of the average position of MLI somas between the basal (0) and distal (1) borders of the Molecular Layer reveals that the Control and Kit KO MLIs have comparable distributions within the molecular layer. Quantification of the thickness of the molecular layer reveals no significant difference between Control and Kit KO conditions. p-values were calculated by a two-tailed t-test with Welch’s correction as necessary, except for Distribution which utilized Mann-Whitney. N refers to the number of different animals whose comparable tissues were evaluated.

**Figure S3:**
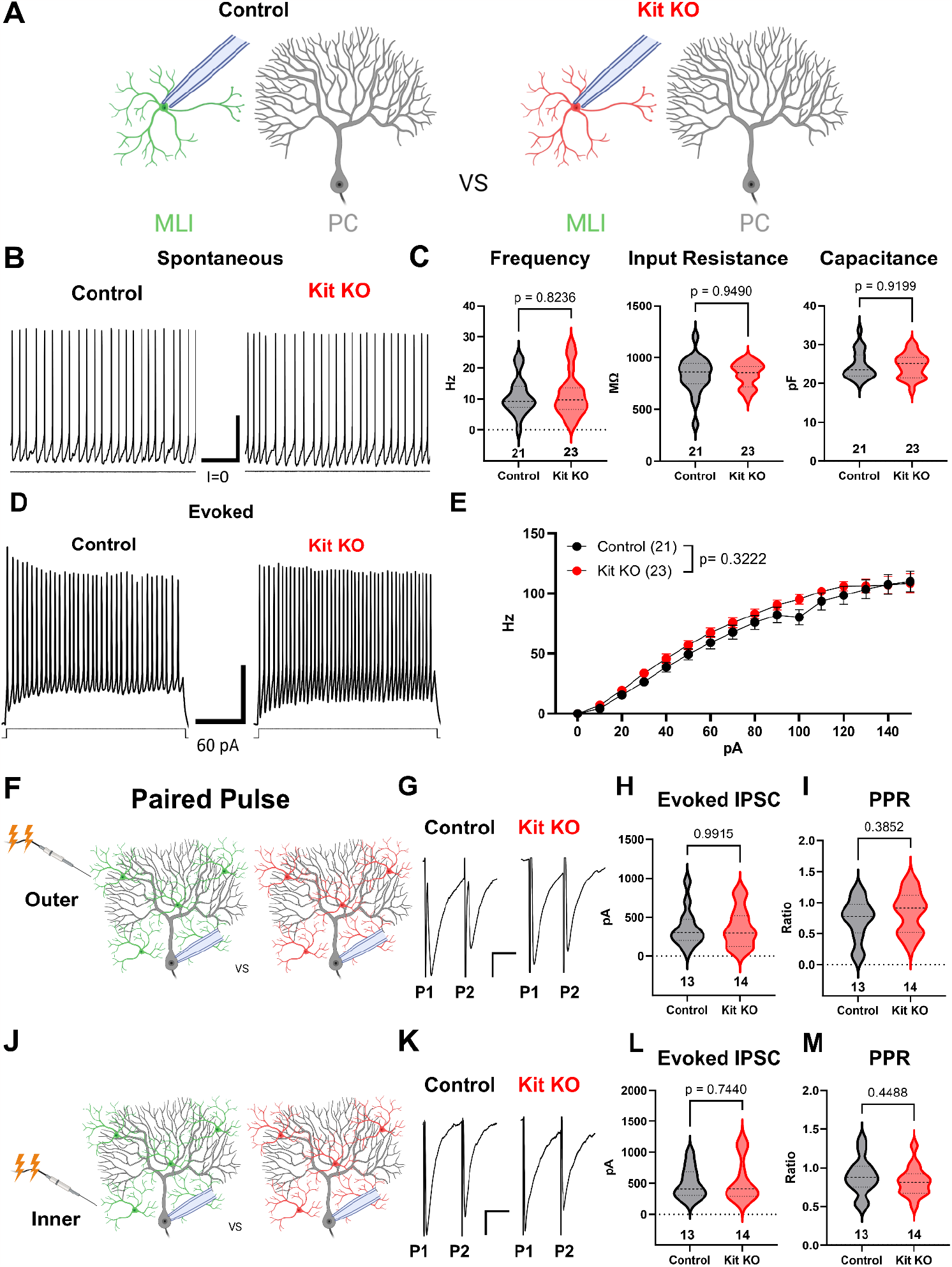
Kit knockout does not alter intrinsic properties of molecular layer interneurons. (**A**) Experimental schema. Patch clamp recordings were performed on Molecular Layer Interneurons in acute cerebellar slice preparations from Control or from Kit KO animals. (**B**,**C**) Patch clamp recordings of Spontaneous action potentials and intrinsic properties in Control and Kit KO MLIs revealed no significant difference in average firing Frequency, membrane Capacitance or Input Resistance. Scale bar 100 ms x 10mV. (**D, E**) Current steps were injected into Control or Kit KO MLIs, example traces (**D**) and quantification (**E**) revealed no significant interaction of genotype and current on evoked mean firing Frequency. Scale bar 100 ms x 10mV (**F, J**) Experimental schema. Paired pulses were delivered via stimulating electrode placed in the Outer or in the Inner molecular layer, and evoked Inhibitory Post Synaptic Currents were evaluated. Example Traces in **G** and **K**. (**H, I; L, M**) The average amplitude of the first evoked inhibitory current, or the Paired Pulse Ratio, did not differ between Control and Kit KO, under either stimulation of the Outer or Inner molecular layer. p-values were calculated by a two-tailed t-test with Welch’s correction as necessary. n refers to the number of different recorded cells per condition.

**Figure S4:**
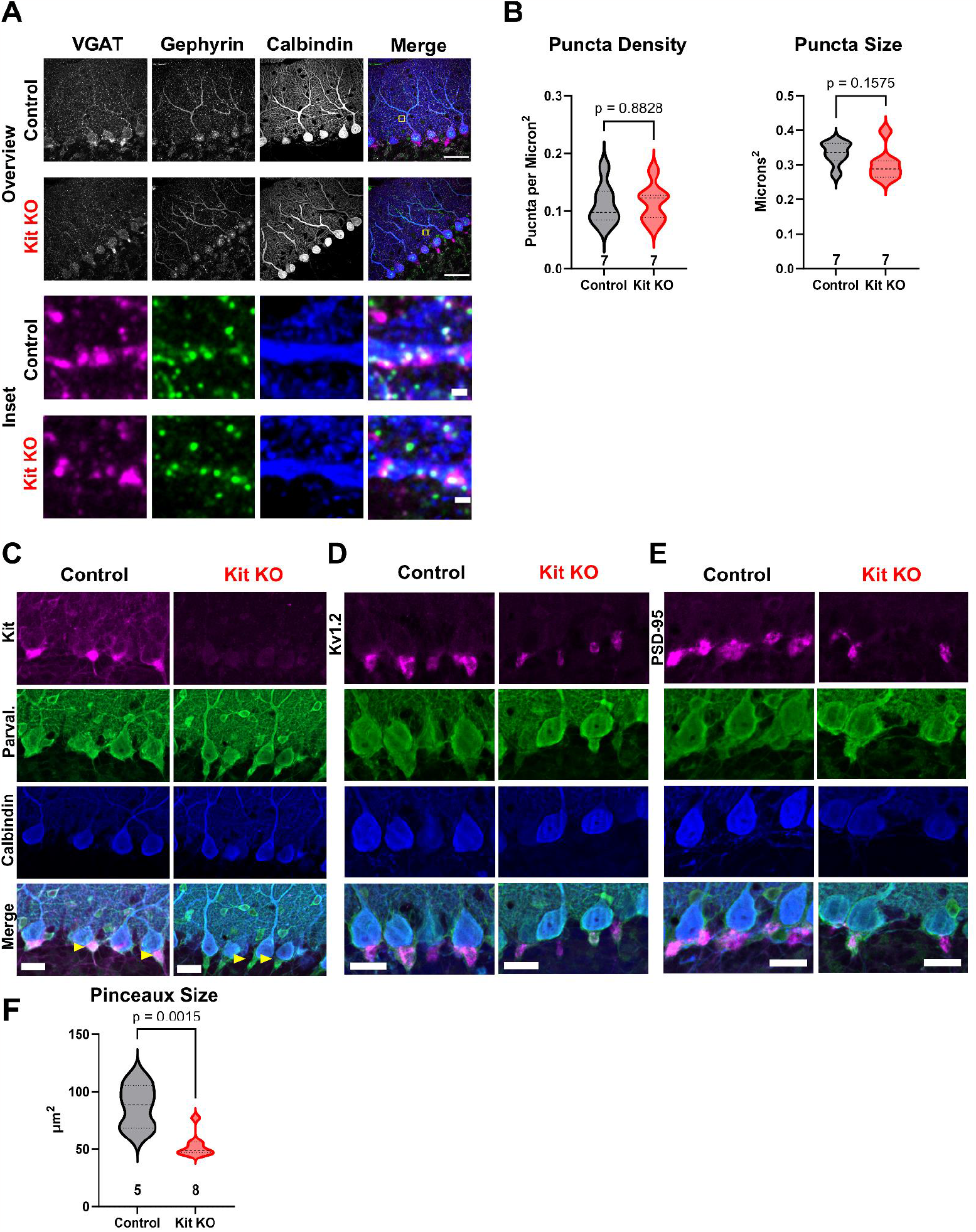
The impact of Kit knockout on the size of molecular layer interneuron synaptic structures. **(A)**Immunohistochemistry and confocal microscopy of presynaptic (VGAT) and postsynaptic (Gephyrin) markers of GABAergic synapses onto Purkinje Cells (Calbindin), for both Control and Kit KO. An Inset from Control (Yellow Box, first Row Merge ROI), demonstrates example triple positive puncta. First two rows, scale bar 50 microns. **(B)**Neither the average density nor the average size (per animal) of GABAergic synaptic puncta onto Purkinje Cells was significantly decreased by Kit KO. (**C, D, E**) Immunohistochemistry and confocal microscopy of markers of the pinceau formation (of MLI axons onto PC soma and initial axon segments). For both Control and Kit KO, Calbindin and Parvalbumin immunoreactivity was determined in conjunction with pinceau markers Kit, Kv1.2, or PSD-95. As evidenced by Parvalbumin positive Calbindin negative pinceau structures in Kit KO in C, pinceau structures do still exist in Kit KO, though they appear smaller. This was affirmed by reduced area of Kv1.2 immunoreactivity, and by PSD-95 immunoreactivity, the latter of which is quantified in (**F**). Scale bars are 50 microns. p-values were calculated by a two-tailed t-test with Welch’s correction as necessary. n refers to the number of different animals per condition.

**Figure S5:**
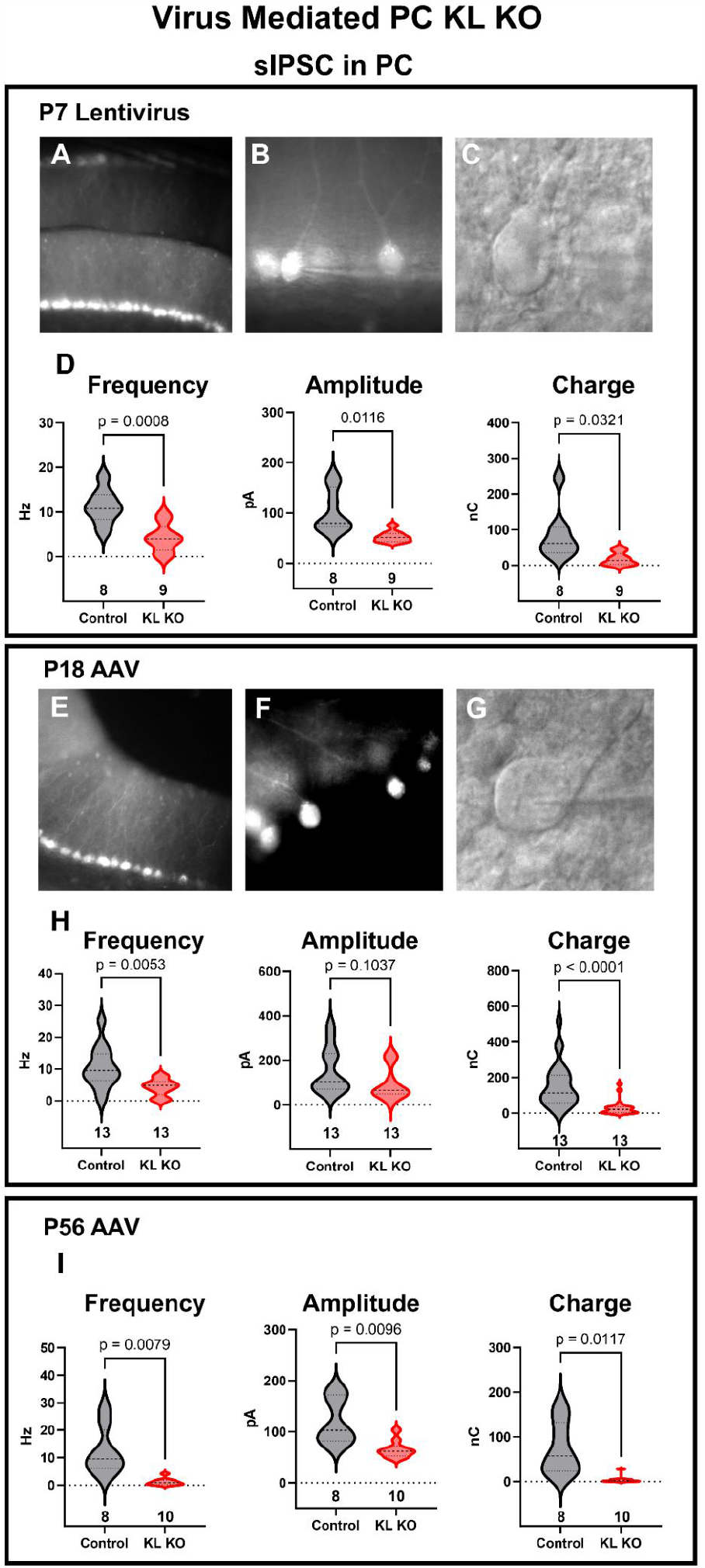
Sparse Acute Depletion of Kit Ligand Reduces GABAergic Input To Purkinje Cells. **(A, B, C**) Lentivirus encoding mCherry-T2A-Cre under the Purkinje Cells specific promoter L7.6 was injected into postnatal day 7 mice homozygous for a Kit Ligand (KL) floxed allele at P7. Demonstrated in **(A)** is an area of direct hit and high transduction. (**B**) Areas with sparse PC transduction were used to record from KL KO or from uninfected adjacent Control PCs, with an example of an IR-DIC identified and patched PC (**C**). (**D**). Analysis of the sIPSCs recorded from P7 Control or sparse KL KO PCs revealed that KL KO reduced both the mean frequency (p=0.0008) and amplitude (p=0.012) and the total inhibitory Charge transfer of GABAergic inhibitory currents in PCs by ∼50%. (**E, F, G**) AAV encoding Cre under the L7.6 promoter was co-injected with an AAV encoding Cre-On mCherry under the Ef1α promoter into the cerebellum of postnatal day 18 or 56 KL floxed homozygous mice. Demonstrated in (**E**) is a direct hit with high transduction, areas as in (**F**) with sparse transduction were selected for recordings of mCherry positive KL KO PCs and adjacent uninfected Control PCs patched under IR-DIC (**G**). (**H, I**). Analysis of the sIPSCs in Control and KL KO PCs at either P18 (**H**) or at P56 (**I**) revealed that mean sIPSC Frequency and Amplitude and total inhibitory Charge transferred were all markedly reduced by postnatal PC KL KO. p-values were calculated by a two-tailed t-test with Welch’s correction as necessary. N in columns refers to the number of different cells recorded.

**Figure S6.**
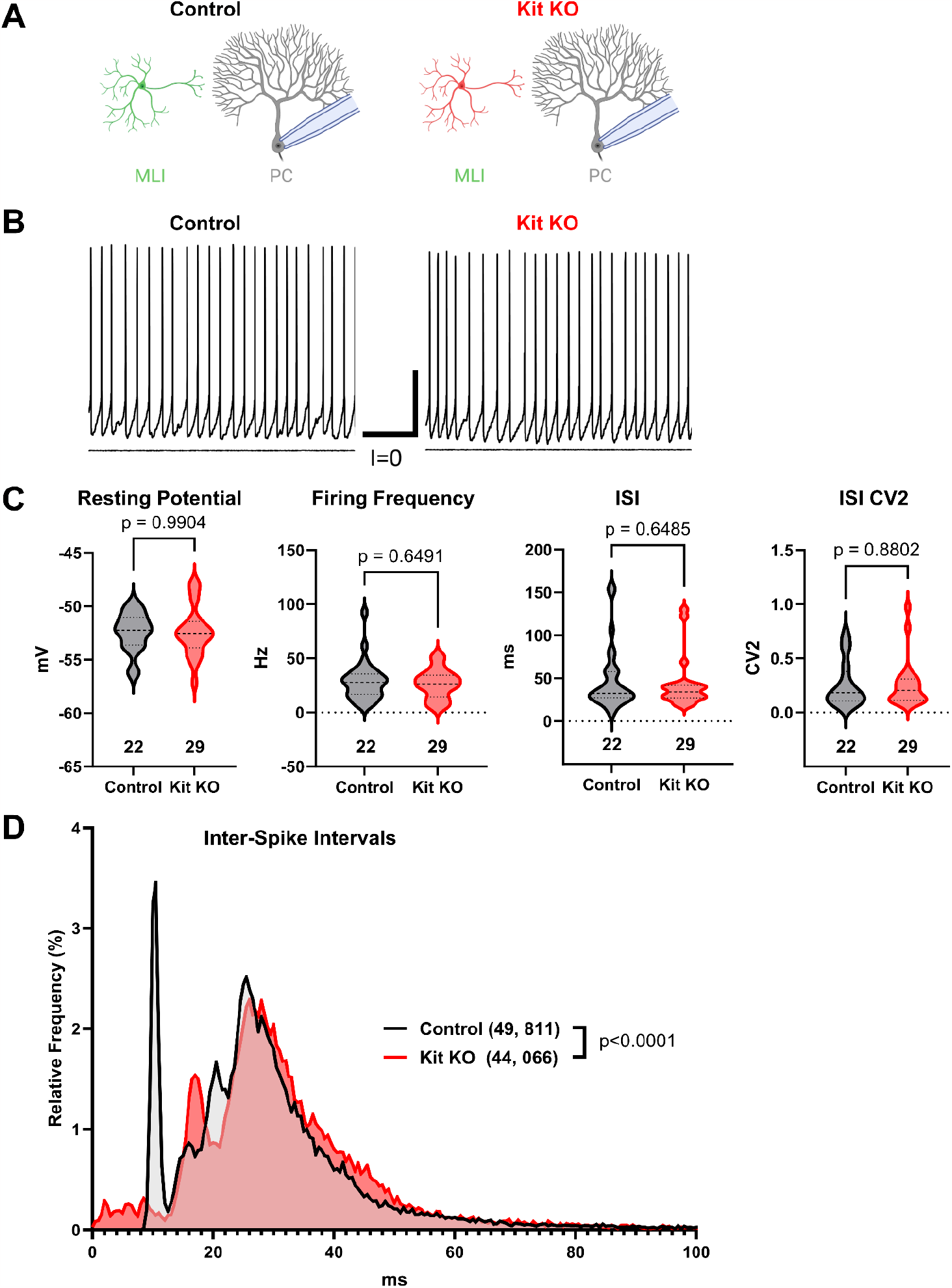
Kit KO does not impact basal Purkinje Cell firing. **A**) Experimental schema and **B**) example traces. Whole cell patch clamp recordings were made from Purkinje cells in acute cerebellar slices from Control or from Kit KO mice. Scale 250 ms x 20 mV. **(C)**Analysis of PC resting membrane potential, spontaneous action potential firing frequency, average per cell Inter-Spike Interval, or per cell ISI CV2, revealed no significant differences between conditions. p values calculated by two-tailed t-test with Welch’s correction as needed. **(D)**A histogram and KS test of individual inter-spike intervals illustrates some differences in the distribution of PC spike timing in Kit KO vs Control conditions.

